# Chasing the Bigfoot: shared molecular patterns among pMHC-I differentiate self from non self peptides

**DOI:** 10.1101/2025.09.08.674516

**Authors:** Etiele S. Silveira, Bruna B. Vian, Alice B. Vian, Eduardo C. Antonio, Renata F. Tarabini, Gustavo F. Vieira

## Abstract

The ability of T cells to discriminate self from non-self peptides is a cornerstone of adaptive immunity, yet the structural principles guiding this process remain poorly defined. Here, we present a proof-of-concept study using physicochemical descriptors derived from images of peptide–MHC (pMHC) surfaces to explore patterns of immunogenicity. We modeled 353 pMHC complexes restricted to HLA-A*02:01, comprising 106 self-derived peptides from the HLA Ligand Atlas and 247 viral epitopes from the Immune Epitope Database. Electrostatic potentials of TCR-facing interfaces were computed and transformed into quantitative features across 92 immunologically relevant regions of interest. Unsupervised hierarchical clustering uncovered subsets of pMHCs forming self-only, viral-only, and mixed clusters, revealing conserved structural fingerprints linked to tolerance or immunogenicity. Heatmap analysis identified close viral–self proximities, highlighting candidate pairs potentially involved in molecular mimicry and autoimmune triggers. Building on these findings, we trained an XGBoost classifier that achieved strong performance (F1 score 0.87; AUC 0.84) in distinguishing self from viral peptides. To our knowledge, this is the first demonstration that structural and physicochemical fingerprints of pMHC complexes are sufficient to discriminate self from non-self. These results establish a framework for computational immunology and provide hypotheses for validation, with implications for autoimmunity, vaccine design, and immunotherapy.

## Introduction

The identification of exogenous proteins is a fundamental step in the proper functioning of adaptive immune responses. To support this, vertebrates not only rely on innate immune defenses but also on a specialized cell-mediated system. In this system, peptide fragments are presented by major histocompatibility complex (MHC) molecules to T lymphocytes, which recognize these peptide-MHC (pMHC) complexes via their specific T cell receptors (TCRs). A key mechanism enabling this specificity is VDJ recombination, a somatic gene rearrangement process that occurs during T cell development. This mechanism generates extensive diversity in the complementarity-determining regions (CDRs) of TCRs, particularly in the CDR3 loops, which are central to the recognition of pMHC complexes and the establishment of interactions with antigen-presenting cells [1]. During development in primary lymphoid organs, T cells undergo a rigorous selection process to eliminate those with high-affinity interactions with self-peptides. This negative selection mechanism induces apoptosis in autoreactive T cells, thereby preventing most of the autoimmune responses. Conversely, T cells that do not strongly bind self-antigens either migrate to secondary lymphoid organs or differentiate into regulatory T cells, which are essential for modulating immune responses - either by suppressing autoreactive T cells or by supporting immune activation under specific conditions [2].

The functional shaping of T cells extends well beyond thymic selection. Throughout life, continuous exposure to environmental antigens and recurrent encounters with pathogens progressively refine the T cell repertoire, enhancing its specificity and adaptability. According to the hygiene hypothesis, the maturation of the immune system is profoundly influenced by microbial exposure during early developmental windows, particularly in childhood, when immune plasticity is at its peak. This concept is further supported by evidence from studies on monozygotic twins, which demonstrate that, despite genetic identity, differential pathogen exposure can result in markedly distinct immune response profiles. Thus, the diversity of antigenic experience during development plays a crucial role in immune system training and long-term homeostasis [3]. In this context, insufficient microbial exposure may blur the boundaries between self and non-self, weakening the immune system’s ability to establish reliable thresholds for cross-reactive responses.

In an effort to better understand how certain peptides become immunogenic, previous studies have investigated sequence-based characteristics such as anchor residues critical for MHC binding, and physicochemical properties like electrostatic potentials, which may influence TCR recognition [4, 5, 6]. Additionally, sequence similarity among epitopes (consensus regions) has been explored as a basis for immune cross-reactivity, in which structurally unrelated pathogens might share epitopes that trigger similar immune responses [7].

Recent findings in cross-reactivity have revealed that structural and physicochemical similarities among peptides from distinct viral sources can facilitate immune recognition even in the absence of significant sequence homology [6, 8]. Expanding upon this concept, our study explores the surface distribution of electrostatic potential patterns in pMHC complexes. We propose that immunogenic pMHCs share molecular signatures and may play a key role in initiating T cell responses. These features could act as fingerprints, informing immunogenicity to TCRs even when sequence identity is minimal, widening immune activation via cross-reactive mechanisms. This supports the hypothesis that immunogenicity is influenced not only by linear peptide sequence, but also by conserved physicochemical elements that guide TCR engagement across diverse antigenic contexts.

To explore this hypothesis, our study began with a clustering analysis based on visual information from TCR-pMHC interaction regions extracted from structural models. Using these image-derived datasets, we identified physicochemical patterns associated with immune response activation. We analyzed pMHC complexes formed by peptides sourced from both immunogenic and healthy tissue databases. Based on the information extracted from the resulting clusters, we trained a supervised classification model. This classifier was then tested on a small, independent set of candidate pMHCs to evaluate its capacity to distinguish between self and non-self peptides based solely on surface properties. This proof of concept offers insight into how structural features could inform immune discrimination, even in the absence of explicit sequence similarity.

## Methods

### Generation of pMHC Models and Image Construction

A total of 106 peptides derived from self-proteins were recovered; 65 peptides from ubiquitously expressed proteins were selected from the Ligand Atlas https://hla-ligand-atlas.org/, filtered for unique HLA-A*02:01 restriction, 9-aa length, presence in ≥3 tissues, and absence of reported immunogenicity in the Immune Epitope Database (IEDB) https://www.iedb.org/, and 41 peptides from cytosolic proteins gelsolin (UniprotID: P06396), actin (UniprotID: P63261), and insulin (UniprotID: P01308). Additionally, 247 immunogenic viral epitopes were retrieved from the IEDB. The peptide sequences obtained from these databases were used to perform molecular docking with HLA-A*02:01 using the DockTope tool [9]. The docked models were subsequently used to compute the electrostatic potentials of the pMHC complex surfaces, specifically in the peptide-binding cleft region. These electrostatic potentials were calculated using DelPhi [10], a well-established tool for predicting electrostatic interactions in biomolecules. The molecular surface visualizations were generated using GRASP2 [11].

The computational time per model varied between 2–3 hours depending on the amino acid composition of the peptide, which directly influenced docking convergence and energy landscape exploration. Each job was executed on a dedicated workstation configured with 8 processing cores and 12 GB of RAM running AutoDock Vina. For the present dataset of 353 complexes, this corresponded to ~700–1,050 computing hours. Although the pipeline for electrostatic potential calculation was partially automated, each complex required manual inspection to confirm correct peptide positioning and epitope representation prior to downstream analyses, ensuring dataset reliability.

### Dataset Clustering

In order to investigate potential distinguishing patterns in the electrostatic potential distributions of pMHC molecular surfaces, an unsupervised exploratory analysis based on hierarchical clustering methods was performed. Initially, predefined regions of interest (ROIs) were extracted from the molecular surface images to focus the analysis on areas of greater functional relevance. As defined by Marcus *et al*., the used ROIs correspond to the contact sites between T cell receptor (TCR) loops and peptide-HLA-A*02:01 (pMHC) complexes. A total of 92 ROI sets were delineated based on this immunological criteria (Supporting Figure S1). This procedure allowed a detailed evaluation of the electrostatic and physicochemical characteristics present at the TCR–pMHC interaction interfaces. The images were processed using the OpenCV library [12] through custom Python scripts. From each ROI, a combination of features were extracted to quantitatively assess color variation within the defined regions of interest (ROIs). We computed the mean and standard deviation of pixel intensities for each ROI across the red, green, and blue channels. These values were used as input for hierarchical clustering, employing the *complete linkage* method with Euclidean distance metrics, resulting in the construction of a dendrogram representing the similarity relationships among the analyzed complexes.

### Predictive Model to differentiate Self from Non-self

The entire dataset of pMHC complexes was also used to train a predictive model using the XGBoost algorithm [13].XGBoost (Extreme Gradient Boosting) is a scalable and efficient machine learning method based on an ensemble of decision trees, optimized through gradient boosting techniques. It is particularly well-suited for structured data, such as the quantitative image features extracted in this study. Feature extraction for each region of interest (ROI) followed the same procedure described in the Dataset Clustering section.

Principal Component Analysis (PCA) was applied to the extracted variables to reduce dimensionality and highlight the features most relevant for distinguishing between classes (0 = self; 1 = non-self). A series of supervised classification models was then trained and validated using increasing numbers of principal components as input features. Because the dataset is imbalanced, with a larger proportion of non-self than self peptides, model selection was primarily guided by F1 score performance, as this metric balances precision and recall and mitigates bias toward the majority class. To ensure robustness, the dataset was split into an independent test set (20% of the data) reserved for final evaluation, while the remaining 80% was further subdivided into training and validation subsets (approximately 64%/16%). Additionally, SHAP analysis was conducted to interpret model predictions and quantify the relative contribution of individual components.

### Heatmap Generation

To visualize pairwise structural/physicochemical proximity, we used the same feature matrix and ROI scheme employed in the HCA (92 TCR–pMHC interface ROIs; per-ROI mean and standard deviation of RGB pixel intensities extracted with OpenCV and custom Python scripts; see Methods). Using this matrix, we computed all pairwise distances with pdist (SciPy, Euclidean metric), and converted the result to a square distance matrix with squareform. Labels for both human and viral proteins were maintained consistently with those used in the hierarchical clustering analysis (HCA), allowing cross-reference between the heatmap and clustering results. Based on literature examples of cross-reactive pMHC pairs (putative molecular mimicry), we defined a distance cutoff of 550 to flag closest matches; this threshold was used only for annotation/highlighting of candidate pairs and not to recompute clustering. The heatmap was rendered in Seaborn (built on Matplotlib), displaying the full inter-complex distance matrix with overlaid markers for pairs below the cutoff. This procedure provides a hypothesis-generating view of viral–self proximity within TCR-facing surfaces, from which candidate mimicry pairs can be prioritized for downstream validation.

## Results

### Clustering uncovers distinct self, viral, and mixed structural sub-patterns

A hierarchical clustering analysis was conducted using the entire dataset of modeled pMHCs, encompassing both self-derived and viral peptides (Supporting Figures S2 and S3). In the initial clustering analysis of the 353 pMHC complexes, no clear large-scale separation between self-derived (human) and non-self (viral) peptides was observed. This outcome was already anticipated, since these two classes are not expected to form unique and mutually exclusive patterns, but rather a spectrum of heterogeneous sub-patterns. Nevertheless, the analysis revealed specific blocks containing pMHC complexes derived from human or viral proteins, indicating that even in the absence of a global dichotomous segregation, certain clusters capture structural proximities and shared physicochemical features among subsets of human-derived peptides.

For example, we identified several clusters composed almost exclusively of self-derived peptides. For clarity, we refer to them here as clusters H1 to H4. Two outgroup clusters (H1 and H2) were observed at the top of the dendrogram, containing two (H2) and three peptides (H1), respectively, all eluted from the Ligand Atlas, which also formed distinct self-only groupings. Cluster H3 encompasses a region also consisting of representatives from eluted peptides derived from proteins with diverse cellular roles, such as MYH11, a smooth muscle myosin heavy chain involved in cytoskeletal contractility, GALC, a lysosomal hydrolase essential for lipid catabolism, DHCR24, an endoplasmic reticulum enzyme in cholesterol biosynthesis, and FAM98A, an RNA-binding protein with regulatory functions—again forming a self-only group. Further down, Cluster H4, located at the central region of the dendrogram, forms a large branch enriched with self-derived peptides, including H2A1, a core histone involved in chromatin organization, Gelsolin (with four distinct peptide representatives), a key actin-binding protein that regulates cytoskeletal remodeling, and BPIFA1, a secreted protein linked to airway mucosal defense—together reinforcing the existence of pure clusters of self-peptides. Other examples similar to those described above can also be found observing the cladogram (Supporting Figures S2 and S3), although they are less conspicuous. Building on previous studies, these observations can be interpreted in light of evidence showing a selective bias in antigen presentation, where only a fraction of genes — and, within those genes, only specific regions — consistently generate MAPs (MHC-associated peptides)[14]. Such patterns, as described in the literature, support the concept that MHC alleles are tuned to preferentially present specific self-regions carrying conserved biochemical signatures indicative of “self,” thereby contributing to peripheral tolerance and the maintenance of physiological homeostasis. Complementary findings from viral proteome analyses have shown that pathogen-derived nonamers structurally and physicochemically more distant from the human self-repertoire tend to exhibit higher predicted TAP translocation efficiencies and stronger MHC binding affinities [15]. These results highlight the evolutionary shaping of multiple steps in the antigen processing and presentation pathway to optimize the immunogenic visibility of non-self regions during infection. Furthermore, previous analyses of electrostatic properties have demonstrated that the mean pairwise similarity among human-derived peptides is higher than that observed between human–viral peptide pairs, reinforcing the concept of a tighter electrostatic clustering of self-ligands and the biophysical cues underlying self-recognition [16].

In parallel to these self-rich clusters, we identified four exclusively viral clusters, designated V1 to V4, each comprising peptides derived from distinct viral families. Cluster V1 contains representatives from hepatitis B virus (HBV), human herpesvirus (HHV), hepatitis C virus (HCV), and influenza A virus (IAV). The remaining clusters (V2 to V4) also include representatives from diverse viral groups (HIV-1, HHV4, HCV, HTLV2, JCV and HPV16), highlighting that their associations cannot be explained by phylogenetic relatedness or by sequence similarity, but rather by the shared physicochemical features of being viral targets and, therefore, potential markers of immunogenicity. The analysis of clusters composed exclusively of viral sequences suggests a recurrent pattern of immunogenicity, in which specific viral regions are consistently recognized by the human immune system. This observation is supported by the presence of public T cell receptors (TCRs), which show a higher propensity to recognize cross-reactive epitopes [14]. The distribution of these TCRs across the population indicates a selective pressure favoring recombinations capable of recognizing viral sequences that are persistently presented to the immune system. Furthermore, the very existence of cross-reactivity among viral peptides provides strong evidence for the presence of conserved structural and physicochemical patterns across distinct viruses — patterns that, if they could be effectively masked or eliminated, would likely have already been targeted by viral evolutionary mechanisms [17]. These findings reinforce the notion that certain viral regions are inherently more immunogenic due to their distinctive characteristics, which differentiate them from the human “self” repertoire, thereby constraining the potential for immune evasion [18]. In addition to the self- and viral-only groups, we also identified clusters displaying more heterogeneous patterns, where self- and non-self–derived peptides were intermixed. Among these, two illustrative examples are clusters M1 and M2. In cluster M1, we identified a closely associated pair consisting of the self-peptide LBR HUMAN ALWNEEALL and the viral peptide HCV ILDSFDPLV. Similarly, cluster M2 revealed two particularly close viral-self pairs: HCV LYPSLIFDI with TMM9B HUMAN LLLAQLSDA, and HHV8 AMLVLLAEI with SCND1 HUMAN QLLAILPEA, which could suggest interesting targets acting as triggers in molecular mimicry evenTaken together, these clustering analyses revealed that, although self- and non-self–derived peptides do not segregate into globally distinct branches, subsets of structural and physicochemical sub-patterns emerge within the dendrogram. It is also important to note that, even though these subgroups emerged as mixed clusters, they may still harbor subtle shared subpatterns that act as markers of immunogenicity or indicators of self-like signatures. Such nuances could, in part, reflect characteristics or biases of the clustering method employed, as seen in studies investigating variants of the same virus that exhibit groups with and without cross-reactivity to a shared pool of lymphocytes [19]. Building upon this observation, we next sought to test whether such sub-patterns could be leveraged to systematically discriminate self from non-self. To this end, we trained a supervised XGBoost classifier using the entire dataset presented in this section, aiming to evaluate its ability to capture and generalize these distinguishing signatures.

### Decoding self versus non-self: a machine learning classifier reveals discriminative molecular features

For the next step, we employed a supervised classification model based on XGBoost, using the same dataset previously analyzed by unsupervised clustering. The input features were extracted from the ROI sets, which provide numerical descriptors that reflect the topography and charge distribution of the original pMHC complexes. With this information, we assessed the model’s ability to discriminate between self-derived and viral peptides.

Since each pMHC complex was represented by 644 numerical descriptors, we applied principal component analysis (PCA) to reduce dimensionality and capture the linear combinations of features that best contribute to variance and separation in the dataset. Initially, we performed PCA on the values extracted from the images (mean and standard deviation of the RGB pixel values). As described in the Methods section, the classifier was tested using the top 5, 10, 15, up to 60 principal components. Classification models trained with increasing numbers of components (5–60) demonstrated consistent performance, with the highest F1 score (0.85) achieved at 55 components. The F1-score values for validation are shown in Fig. 1. Based on these results, the 55-component configuration was selected as the optimal model, with Accuracy of 0.78, Precision of 0.83, and Recall of 0.86, reflecting balanced performance across all metrics. For this approach, the 5-fold validation results are presented in Fig. 2A. From the 20% of peptides set aside for testing, the model achieved a precision of 0.88, recall of 0.86, F1 score of 0.87, and accuracy of 0.82. The confusion matrix (Fig. 2B) showed that only 6 self-peptides were misclassified as viral, and 7 viral peptides were misclassified as self. In addition, the ROC curve (Fig. 2C) yielded an AUC of 0.84, further confirming the discriminative power of the model by quantifying its ability to correctly rank self- and non-self–derived peptides across all classification thresholds. These results highlight the ability of physicochemical image-derived data to characterize the origin of pMHCs and their consequent potential to determine immunogenicity within an immunological context, further underscoring the robustness and reliability of our image-based analyses, as consistently demonstrated in our previous studies [20, 21, 22]. To assess whether data bias influenced the results, we performed label randomization tests using the same optimal PCA-55 configuration identified in the main analysis. In the first test, labels were reassigned for 50% of the dataset, while the remaining half kept their original annotation. This procedure still yielded a F1 score of 0.72, indicating that the correctly labeled subset retained a sufficiently strong signal to support reasonable classification performance. In a second round, labels were randomized for 100% of the dataset. Even under this condition, by chance, a fraction of structures still retained their original labels, allowing us to evaluate the effect of residual signal. In this case, the model achieved a reduced F1 score of 0.58, confirming that the robust performance observed in the main analysis was not an artifact of data bias. Additionally, we conducted a SHAP (Shapley Additive Explanations) analysis to interpret the contribution of each principal component to the classification. This approach allowed us to identify the Principal Components (PCs) with the highest impact on the model’s predictions and, by mapping these components back to the original images, to pinpoint the specific regions within the pMHC interaction interface that carry the most relevant information for group separation. These insights not only enhance the interpretability of our approach but also provide a framework for guiding future studies aiming to refine image-based descriptors of pMHC complexes. The detailed visualizations and supporting analyses are provided in the Supporting Material (Table S1; Figure S4; Figure S5).

**Fig. 1:**
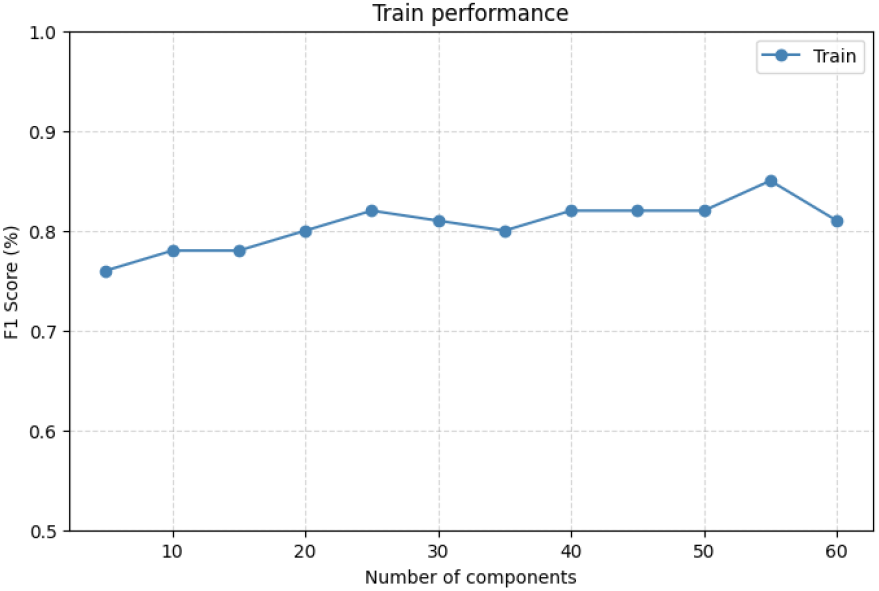
Training performance (F1-Score) as a function of the number of PCA components: Evolution of the F1-Score (%) for varying numbers of PCA components. The model achieves optimized performance with 55 components, suggesting an optimal balance between dimensionality reduction and predictive power.

**Fig. 2:**
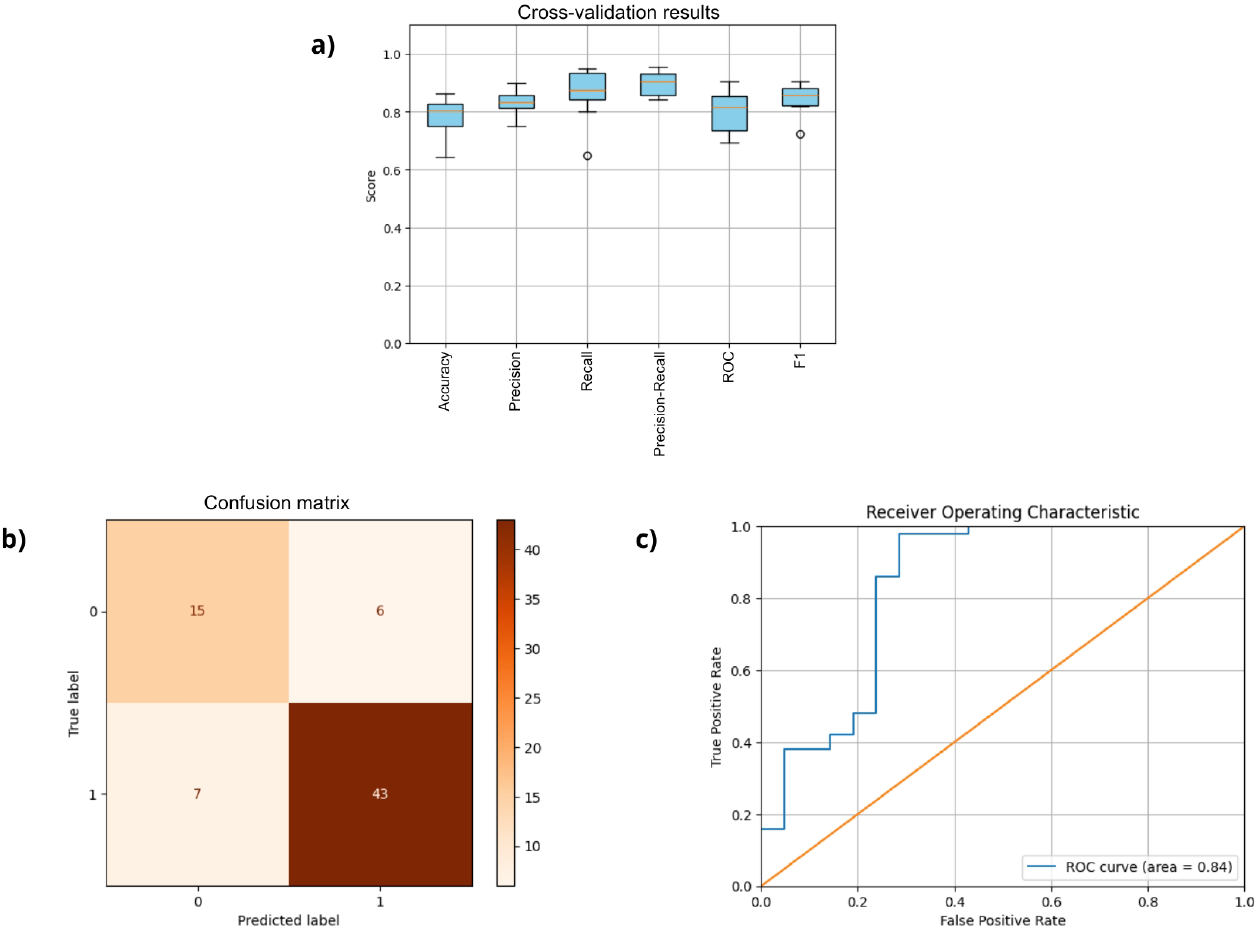
Performance evaluation of the XGBoost classifier using 55 principal components: (a) Cross-validation results showing accuracy, precision, recall, precision-recall curve, ROC curve, and F1-score; (b) Confusion matrix; (c) ROC curve with AUC of 0.84.

It is important to highlight that the likely success of our predictor is supported by the availability of datasets generated from healthy tissues, such as the HLA Ligand Atlas, which provides a valuable baseline of self-peptides. In contrast, databases like the IEDB predominantly contain tumor-derived immunogenic peptides, when we set “human source,” further emphasizing the imbalance in the sources available for training and validation. In fact, most publicly available datasets of human HLA-presented peptides are derived from tumor immunopeptidomics studies, often generated through mass spectrometry analyses of malignant tissues [23]. This creates an inherent bias in the current repositories, as the majority of “human” peptides annotated in these datasets are, in fact, tumor-associated peptides, which tend to be more immunogenic than those derived from healthy tissues [24]. Consequently, this scarcity of immunopeptidomic data from non-malignant samples limits the accurate definition of the true self-peptidome and introduces challenges in understanding peripheral tolerance and the landscape of immunogenicity under physiological conditions.

### Structural correlations between viral and human pMHCs reveal potential zones of molecular mimicry

The heatmap generated from the pairwise distance analysis Fig. 3 highlights regions of close structural similarity between viral and human pMHC complexes, revealing potential zones where molecular mimicry may occur. These proximities, observed even among peptides derived from proteins normally expressed on the surface of antigen-presenting cells, underscore the possibility that certain viral epitopes share key physicochemical and structural features with self-peptides. Such overlaps could act as triggers for autoreactive T-cell responses, particularly in contexts where peripheral tolerance is disrupted. This approach serves as a hypothesis-generating framework, providing a systematic way to identify candidate viral–self-peptide pairs that warrant deeper experimental investigation. By mapping these structural proximities, our analysis offers a starting point to uncover novel links between viral infections and autoimmune manifestations, especially in scenarios where the triggering viral epitope and the targeted self-protein remain unknown.

**Fig. 3:**
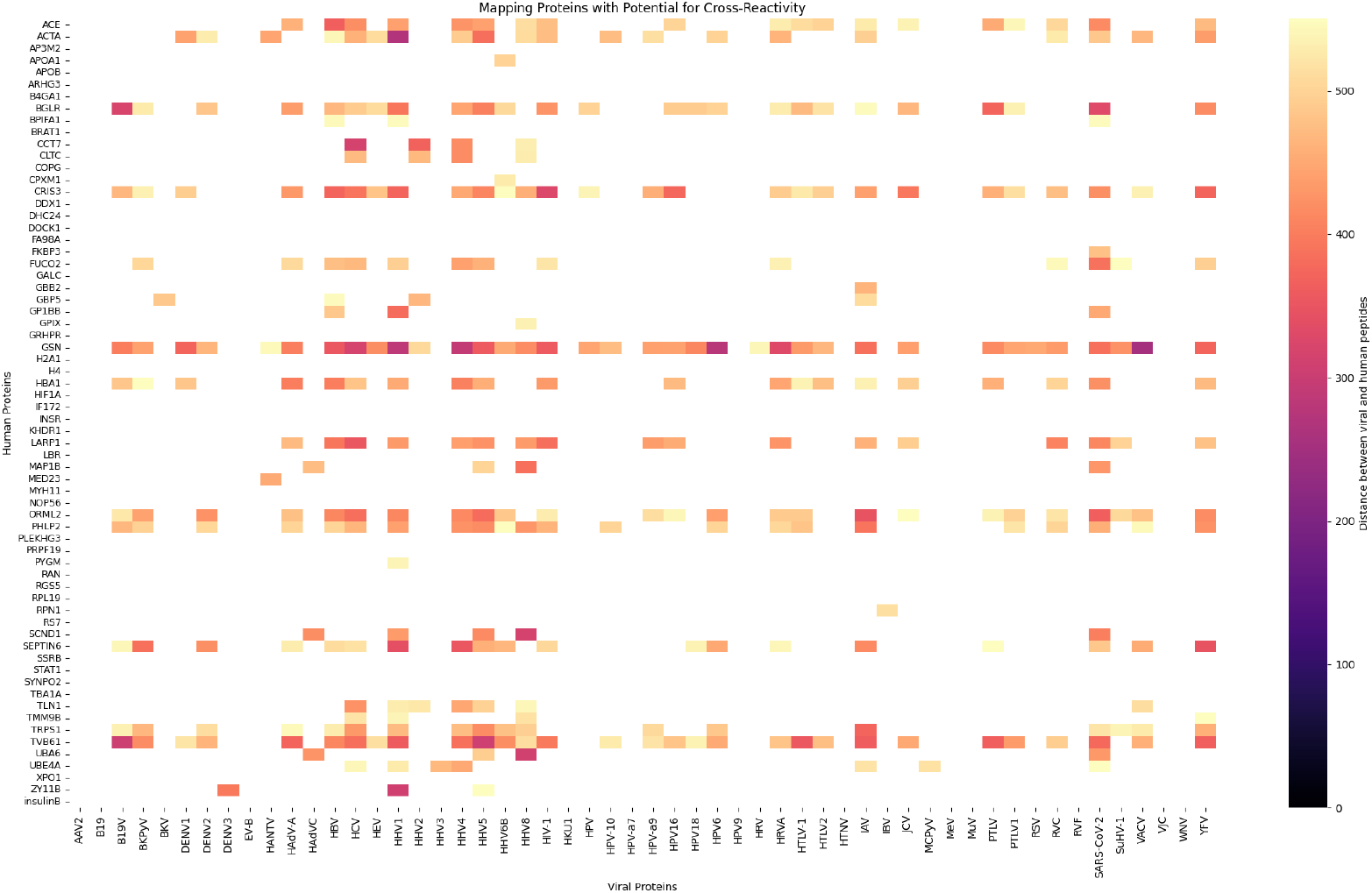
Heatmap showing pairwise distances between peptides derived from human proteins (rows) and viral proteins (columns). Colors represent the distance between peptides, with yellow indicating larger distances (more dissimilar) and black indicating smaller distances (more similar).

Within the heatmap, a distinct block of viral representatives was evident, comprising human adenoviruses, multiple hepatitis viruses, and diverse human herpesviruses, indicating a higher concentration of structural similarities with human-derived peptides. In another region of the heatmap, additional viral representatives such as Influenza A, SARS-CoV-2, and Yellow Fever virus also exhibited several matches with human proteins, although not arranged in family-specific clusters as observed in the upper block. Some of these viruses, including Influenza A [25] and SARS-CoV-2 [26], have already been associated with the onset or exacerbation of autoimmune disorders, further reinforcing their relevance as potential triggers in cross-reactive immune responses.

On the human side, proteins such as ACTB, GSN, TRPS1, TVB61, CRIS3, and BGLR emerged as recurrent counterparts, displaying structural convergence with peptides from multiple viral origins. These patterns highlight not only the diversity of potential viral triggers but also pinpoint human proteins that may act as hubs for cross-reactivity, providing a robust framework for prioritizing candidates in future mechanistic and structural validation studies.

Although the association between molecular mimicry and the onset of autoimmunity is well recognized, there remains a significant paucity of studies that delineate, with molecular precision, the specific T-cell epitopes that act as initiators of pathogenic responses. While several studies have established correlations between particular MHC alleles, pathogen exposure, and autoimmune phenotypes [27, 28], the precise antigenic targets of the T-cell repertoire in these contexts remain poorly characterized [29]. Only a limited number of cases have been comprehensively elucidated, in which a clear structural and functional correspondence between a viral-derived epitope and its self counterpart has been experimentally validated [30, 31]. This persistent gap in the literature underscores the need for systematic frameworks — such as the structural proximity analyses presented here — to refine the identification of candidate viral–self pairs for subsequent experimental interrogation. Such integrative approaches hold the potential to elucidate the critical molecular events underlying the disruption of peripheral tolerance and the progression toward autoimmune pathology.

## Conclusion and Future Directions

In this study, we provide the first evidence that physicochemical image-derived descriptors of pMHC surfaces can reliably discriminate self from non-self peptides. This finding highlights that structural and electrostatic signatures embedded in the TCR-facing interface encode sufficient information to support immunological classification in the absence of sequence identity. By combining unsupervised clustering and a supervised XGBoost classifier, we demonstrate that viral epitopes converge on shared molecular patterns that underlie their recurrent immunogenicity, whereas self-derived peptides cluster within more constrained electrostatic landscapes consistent with tolerance. This unprecedented result fills a critical gap in the field, where previous studies had only correlated sequence-based or allele-level features with immunogenicity. To our knowledge, this is the first demonstration that structural and physicochemical fingerprints extracted from pMHC complexes are sufficient to discriminate self from non-self, opening a new avenue for understanding immune recognition and cross-reactivity beyond sequence-based paradigms.

Beyond establishing a novel computational framework, our analyses also provide actionable hypotheses for experimental follow-up. In particular, the heatmap of viral–self proximities highlights candidate epitopes and human proteins that may serve as starting points for laboratory validation, offering a systematic route to probe potential triggers of autoimmune disturbances. By integrating these image-based descriptors with high-throughput immunopeptidomics and single-cell TCR sequencing, future studies could bridge computational predictions with functional immunology, ultimately refining our understanding of T cell recognition, tolerance, and cross-reactivity, and informing vaccine development and immunotherapeutic strategies.

## References

1. Jan Černý and Igor Stříž. Adaptive innate immunity or innate adaptive immunity? Clinical Science (London), 133(14):1549–1565, 2019. Published 2019 Jul 17.

2. D. Campillo-Davo, D. Flumens, and E. Lion. The quest for the best: How tcr affinity, avidity, and functional avidity affect tcr-engineered t-cell antitumor responses. Cells, 9(7):1720, Jul 2020.

3. C. Tabilas, D. S. Iu, C. W. P. Daly, K. J. Yee Mon Reynaldi, S. P. Wesnak, J. K. Grenier, M. P. Davenport, N. L. Smith, A. Grimson, and B. D. Rudd. Early microbial exposure shapes adult immunity by altering cd8+ t cell development. Proceedings of the National Academy of Sciences of the United States of America, 119(49):e2212548119, Dec 2022. Epub 2022 Nov 28.

4. D.A. Antunes, M.M. Rigo, J.P. Silva, S.P. Cibulski, M. Sinigaglia, J.A. Chies, and G.F. Vieira. Structural in silico analysis of cross-genotype-reactivity among naturally occurring hcv ns3-1073-variants in the context of hla-a*02:01 allele. Molecular Immunology, 48(12–13): 1461–1467, 2011.

5. M.F. Mendes, D.A. Antunes, M.M. Rigo, M. Sinigaglia, and G.F. Vieira. Improved structural method for t-cell cross-reactivity prediction. Molecular Immunology, 67(2 Pt B):303–310, 2015. Epub 2015 Jul 2.

6. D.A. Antunes, M.M. Rigo, M.V. Freitas, M.F.A. Mendes, M. Sinigaglia, G. Lizée, L.E. Kavraki, L.K. Selin, M. Cornberg, and G.F. Vieira. Interpreting t-cell cross-reactivity through structure: Implications for tcr-based cancer immunotherapy. Frontiers in Immunology, 8:1210, 2017. Published online 2017 Oct 4.

7. A. Grifoni, D. Weiskopf, S.I. Ramirez, J. Mateus, J.M. Dan, C.R. Moderbacher, S.A. Rawlings, A. Sutherland, L. Premkumar, R.S. Jadi, D. Marrama, A.M. de Silva, A. Frazier, A.F. Carlin, J.A. Greenbaum, B. Peters, F. Krammer, D.M. Smith, S. Crotty, and A. Sette. Targets of t cell responses to sars-cov-2 coronavirus in humans with covid-19 disease and unexposed individuals. Cell, 181(7):1489–1501.e15, 2020. Published online 2020 May 20.

8. M.F.A. Mendes, M. de Souza Bragatte, P. Vianna, M.V. de Freitas, I. Pöhner, S. Richter, R.C. Wade, F.M. Salzano, and G.F. Vieira. Matchtope: A tool to predict the cross reactivity of peptides complexed with major histocompatibility complex i. Frontiers in Immunology, 13:930590, 2022. Published online 2022 Oct 28.

9. Marcos M. Rigo, Dinler A. Antunes, Maycon Vaz de Freitas, Murilo Fabiano de Almeida Mendes, Luiz Meira, Maristela Sinigaglia, and Gerson F. Vieira. Docktope: a web-based tool for automated pmhc-i modelling. Scientific Reports, 5:18413, 2015.

10. Michael K. Gilson and Barry Honig. Calculation of the total electrostatic energy of a macromolecular system: solvation energies, binding energies, and conformational analysis. Proteins: Structure, Function, and Bioinformatics, 4(1): 7–18, 1988.

11. Donald Petrey and Barry Honig. Grasp2: visualization, surface properties, and electrostatics of macromolecular structures and sequences. Methods in Enzymology, 374: 492–509, 2003.

12. Gary Bradski and Adrian Kaehler. Learning OpenCV: Computer Vision with the OpenCV Library. O’Reilly Media, 2008.

13. Tianqi Chen and Carlos Guestrin. XGBoost: A scalable tree boosting system. In Proceedings of the 22nd ACM SIGKDD International Conference on Knowledge Discovery and Data Mining, KDD ‘16, pages 785–794, New York, NY, USA, 2016. Association for Computing Machinery.

14. Hillary Pearson, Tariq Daouda, Diana Paola Granados, Chantal Durette, Eric Bonneil, Mathieu Courcelles, Anja Rodenbrock, Jean-Philippe Laverdure, Caroline Côté, Sylvie Mader, Sébastien Lemieux, Pierre Thibault, and Claude Perreault. Mhc class i–associated peptides derive from selective regions of the human genome. The Journal of Clinical Investigation, 126(12):4690–4701, 12 2016.

15. Daniele Santoni and Giovanni Felici. An immunological glimpse of human virus peptides: Distance from self, mhc class i binding, proteasome cleveage, tap transport and sequence composition entropy. Virus Research, 317:198814, 2022.

16. Marco Antônio M Pretti, Gustavo Fioravanti Vieira, Mariana Boroni, and Martín H Bonamino. Unveiling cross-reactivity: implications for immune response modulation in cancer. Briefings in Bioinformatics, 26(1):bbaf012, 01 2025.

17. L K Selin, S R Nahill, and R M Welsh. Cross-reactivities in memory cytotoxic t lymphocyte recognition of heterologous viruses. Journal of Experimental Medicine, 179(6):1933–1943, 06 1994.

18. E. G. M. Berkhoff, E. de Wit, M. M. Geelhoed-Mieras, C. M. Boon, J. Symons, R. A. M. Fouchier, A. D. M. E. Osterhaus, and G. F. Rimmelzwaan. Functional constraints of influenza a virus epitopes limit escape from cytotoxic t lymphocytes. Journal of Virology, 79(17): 11239–11246, 2005.

19. Marcus F.A. Mendes, Dinler A. Antunes, Maurício M. Rigo, Marialva Sinigaglia, and Gustavo F. Vieira. Improved structural method for t-cell cross-reactivity prediction. Molecular Immunology, 67(2, Part B):303–310, 2015.

20. Dinler A. Antunes, Maurício M. Rigo, Martiela V. Freitas, Marcus F. A. Mendes, Marialva Sinigaglia, Gregory Lizée, Lydia E. Kavraki, Liisa K. Selin, Markus Cornberg, and Gustavo F. Vieira. Interpreting t-cell cross-reactivity through structure: Implications for tcr-based cancer immunotherapy. Frontiers in Immunology, Volume 8 - 2017, 2017.

21. Dinler A. Antunes, Maurício M. Rigo, Jader P. Silva, Samuel P. Cibulski, Marialva Sinigaglia, José A.B. Chies, and Gustavo F. Vieira. Structural in silico analysis of cross-genotype-reactivity among naturally occurring hcv ns3-1073-variants in the context of hla-a*02:01 allele. Molecular Immunology, 48(12): 1461–1467, 2011.

22. S. Zhang, R. K. Bakshi, P. V. Suneetha, P. Fytili, D. A. Antunes, G. F. Vieira, R. Jacobs, C. S. Klade, M. P. Manns, A. R. M. Kraft, H. Wedemeyer, V. Schlaphoff, and M. Cornberg. Frequency, private specificity, and crossreactivity of preexisting hepatitis c virus (hcv)-specific cd8+ t cells in hcv-seronegative individuals: Implications for vaccine responses. Journal of Virology, 89(20): 11070–11082, 2015.

23. Lena Katharina Freudenmann, Ana Marcu, and Stefan Stevanović. Mapping the tumour human leukocyte antigen (hla) ligandome by mass spectrometry. Immunology, 154(3): 331–345, 2018.

24. C. Ragone, C. Manolio, B. Cavalluzzo, A. Mauriello, M. L. Tornesello, F. M. Buonaguro, et al. Identification and validation of viral antigens sharing sequence and structural homology with tumor-associated antigens (taas). Journal for ImmunoTherapy of Cancer, 9(9):e002694, 2021.

25. Shunyu Xie, Jintian Wei, and Xiaohui Wang. The intersection of influenza infection and autoimmunity. Frontiers in Immunology, Volume 16 - 2025, 2025.

26. C. Sharma and J. Bayry. High risk of autoimmune diseases after COVID-19. Nature Reviews Rheumatology, 19(7):399–400, 2023.

27. Sandra S. Soldan and Paul M. Lieberman. Epstein–barr virus and multiple sclerosis. Nature Reviews Microbiology, 21:51–64, January 2023. Published online 05 August 2022, accepted 27 June 2022.

28. S. Sakaue, S. Gurajala, M. Curtis, et al. Tutorial: a statistical genetics guide to identifying hla alleles driving complex disease. Nature Protocols, 18(9): 2625–2641, 2023.

29. Jörg Christoph Prinz. Immunogenic self-peptides - the great unknowns in autoimmunity: Identifying t-cell epitopes driving the autoimmune response in autoimmune diseases. Frontiers in Immunology, Volume 13 - 2022, 2023.

30. Qin Ouyang, Nathan E. Standifer, Huilian Qin, Peter Gottlieb, C. Bruce Verchere, Gerald T. Nepom, Rusung Tan, and Constadina Panagiotopoulos. Recognition of hla class i–restricted-cell epitopes in type 1 diabetes. Diabetes, 55(11):3068–3074, 11 2006.

31. Kai W. Wucherpfennig. Mechanisms for the induction of autoimmunity by infectious agents. The Journal of Clinical Investigation, 108(8):1097–1104, 10 2001.

