## Supplementary Material for "Chasing the Bigfoot: shared molecular patterns among pMHC-I differentiate self from non self peptides"

### Supporting Material

#### Overview

This document provides supporting material for the manuscript entitled "Chasing the Bigfoot: shared molecular patterns among pMHC-I differentiate self from non self peptides".

#### Supporting Tables

**Table S1:** Top 10 most important principal components (PCs) in the XGBoost model using PCA 55, showing feature importance (%) and cumulative contribution (%).

| PC | Feature Importance (%) | Cumulative (%) |
|---|---|---|
| PC12 | 11.1966 | 11.1966 |
| PC23 | 7.7992 | 18.9958 |
| PC5 | 7.4898 | 26.4856 |
| PC28 | 5.4533 | 31.9388 |
| PC17 | 5.4237 | 37.3626 |
| PC46 | 4.5704 | 41.9330 |
| PC21 | 3.6773 | 45.6103 |
| PC7 | 3.1008 | 48.7111 |
| PC2 | 2.7495 | 51.4606 |
| PC4 | 2.4657 | 53.9263 |

#### Supporting Figures

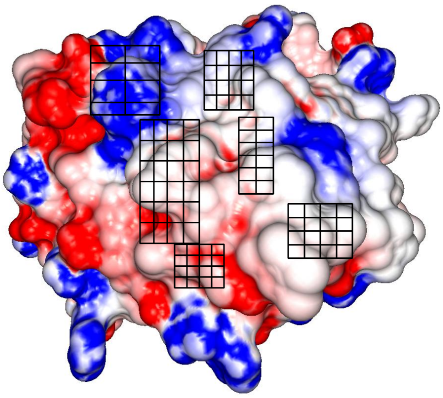

**Figure S1:** Electrostatic potential map of HLA-A02:01 with the 92 ROIs highlighted, complexed with the LIDQYLYYL peptide.

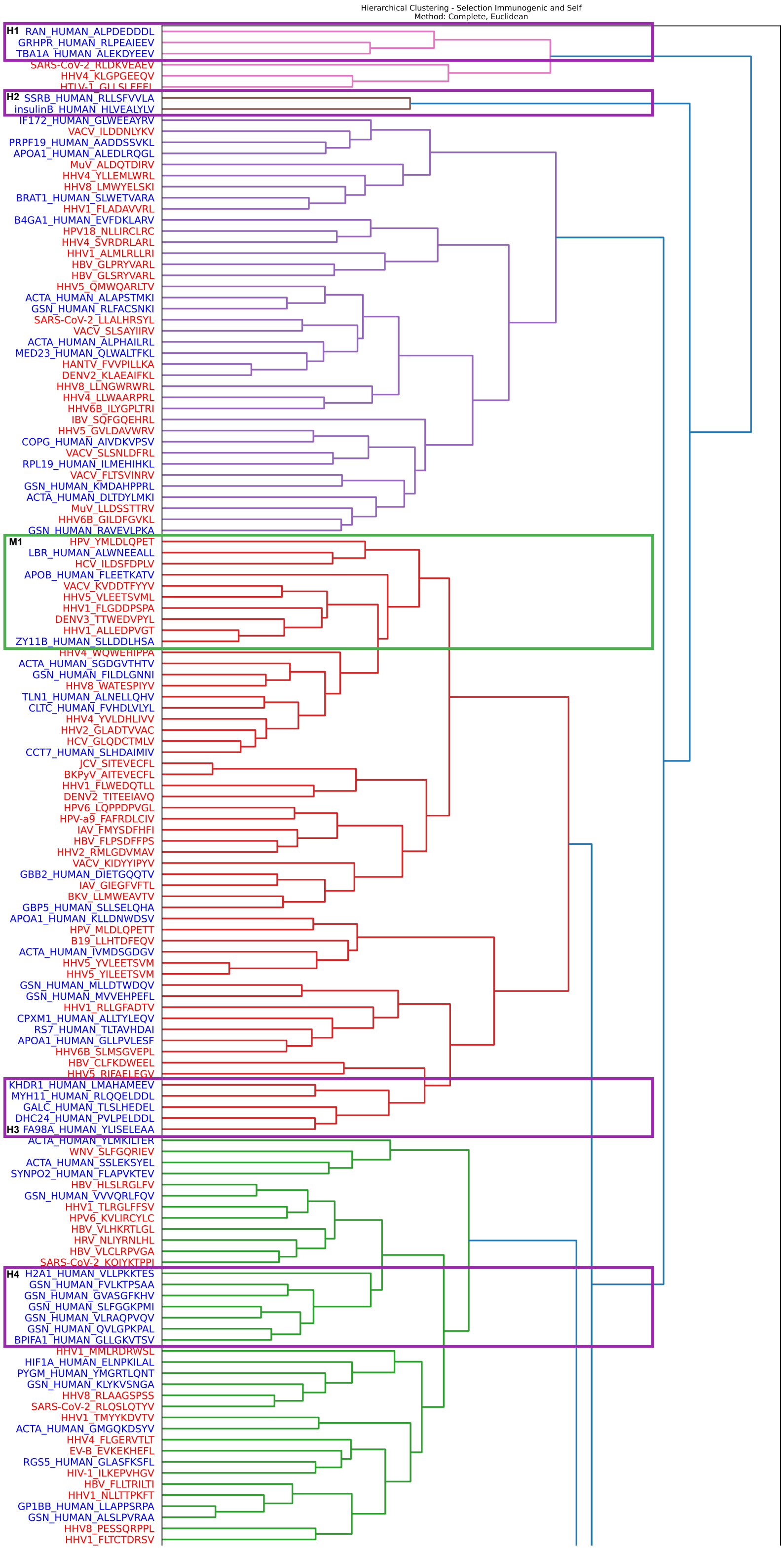


**Figure S2: Hierarchical clustering of peptide–MHC complexes derived from viral and human proteins(top part).** The dataset comprises peptides originating from viral proteins (labels in red) and human self-proteins (labels in blue). Purple rectangles highlight clusters exclusively composed of human-derived peptides, green rectangles indicate mixed clusters containing both viral and self peptides, and orange rectangles denote clusters formed solely by viral-derived peptides.

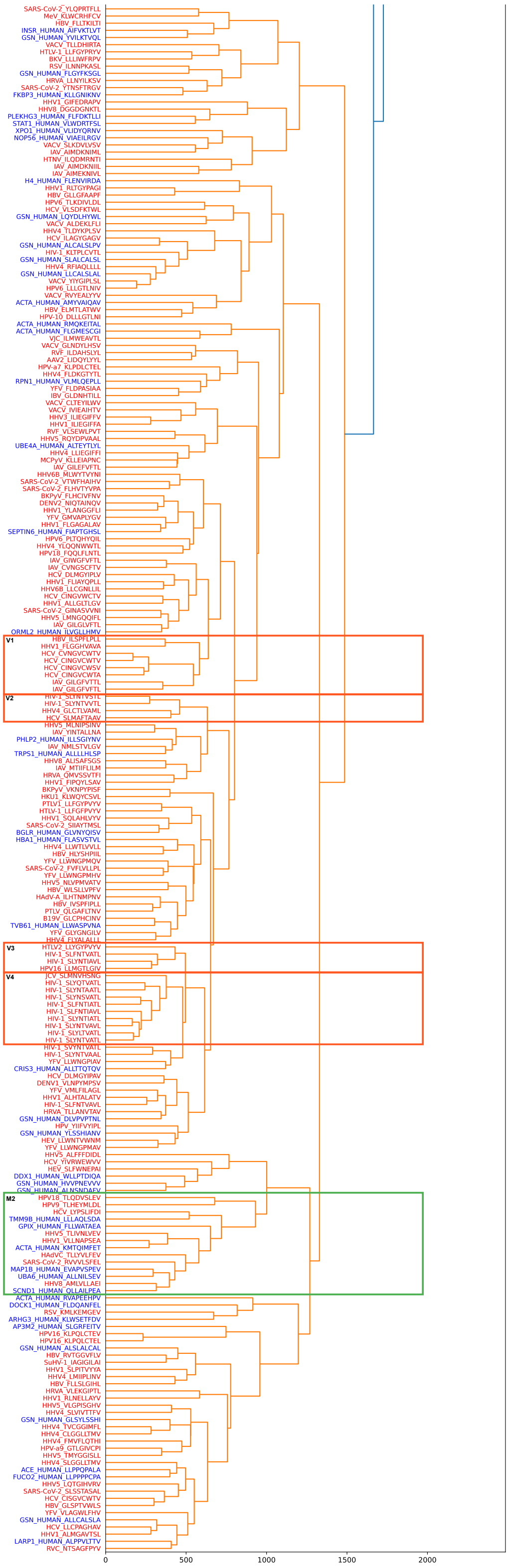


**Figure S3: Hierarchical clustering (bottom part).**

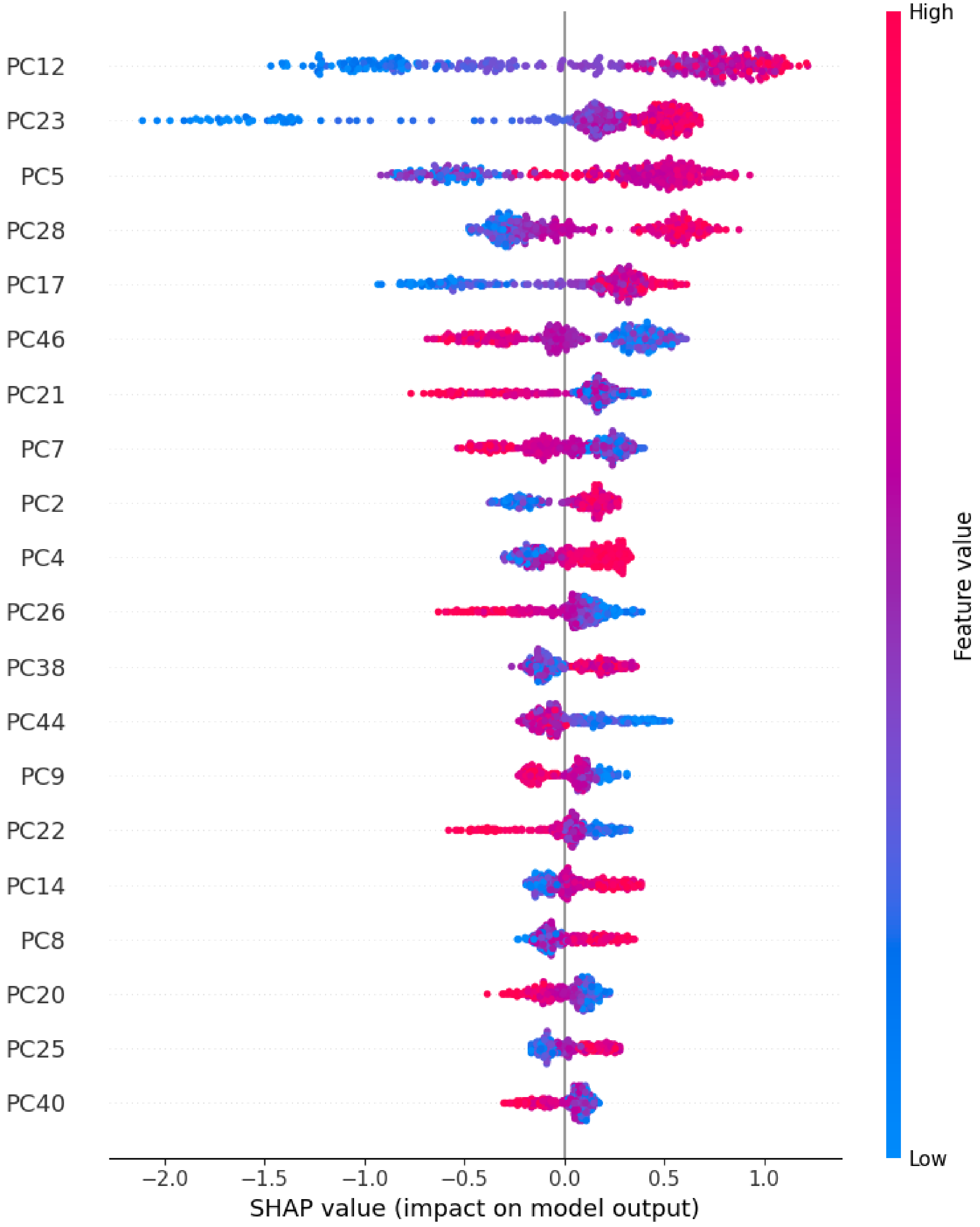


**Figure S4: SHAP analysis of XGBoost model using 55 principal components (PCA-55)**

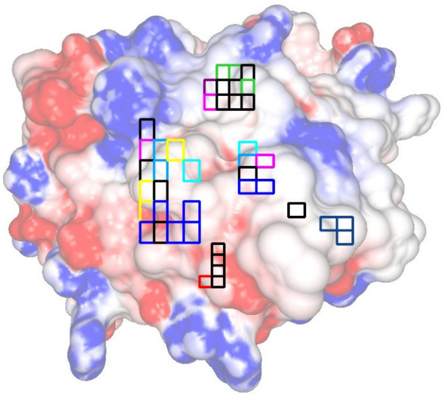

**Figure S5:** Electrostatic potential of HLA-A02:01 complexed with the peptide LIDQYLYYL, annotated with the SHAP contribution of the ten most important principal components (PCs) from the best-performing XGBoost model. Colored squares indicate the regions associated with each PC: PC12 (blue), PC23 (green), PC5 (red), PC28 (cyan), PC17 (magenta), PC46 (yellow), PC21 (orange), PC7 (purple), PC2 (brown), and PC4 (lime green). Black squares represent overlapping contributions between PCs.
